# A broad-spectrum α-glucosidase of glycoside hydrolase family 13 from *Marinovum* sp., a member of the *Roseobacter* clade

**DOI:** 10.1101/2023.03.31.535182

**Authors:** Jinling Li, Janice W.-Y. Mui, Bruna M. da Silva, Douglas E.V. Pires, David B. Ascher, Niccolay Madiedo Soler, Ethan D. Goddard-Borger, Spencer J. Williams

## Abstract

Glycoside hydrolases (GHs) are a diverse group of enzymes that catalyze the hydrolysis of glycosidic bonds. The Carbohydrate-Active enZymes (CAZy) classification organizes GHs into families based on sequence data and function, with fewer than 1% of the predicted proteins characterized biochemically. Consideration of genomic context can provide clues to infer possible enzyme activities for proteins of unknown function. We used the MultiGeneBLAST tool to discover a gene cluster in *Marinovum* sp., a member of the marine *Roseobacter* clade, that encodes homologues of enzymes belonging to the sulfoquinovose monooxygenase pathway for sulfosugar catabolism. This cluster lacks a gene encoding a classical family GH31 sulfoquinovosidase candidate, but which instead includes an uncharacterized family GH13 protein (*Ms*GH13) that we hypothesized could be a non-classical sulfoquinovosidase. Surprisingly, recombinant *Ms*GH13 lacks sulfoquinovosidase activity and is a broad spectrum α-glucosidase that is active on a diverse array of α-linked disaccharides, including: maltose, sucrose, nigerose, trehalose, isomaltose, and kojibiose. Using AlphaFold, a 3D model for the *Ms*GH13 enzyme was constructed that predicted its active site shared close similarity with an α-glucosidase from *Halomonas* sp. H11 of the same GH13 subfamily that shows narrower substrate specificity.

## Introduction

Glycoside hydrolases (GHs), which catalyze the hydrolysis of glycosidic bonds, are a large and diverse group of enzymes. GHs have been classified into families in the Carbohydrate-Active enZymes (CAZy) database (www.cazy.org1 and www.cazypedia.org2). The CAZy database, which consists of around 170 families, organizes GHs into families based on sequence data and function. However, fewer than 1% of the predicted proteins within the CAZy database have been biochemically characterized, and therefore sequence-based predictions of functions are the primary means for assigning possible functions to uncharacterized GH proteins. Often, especially within bacteria, genes encoding specific GHs occur within gene clusters that contain other genes encoding related enzymes that enable the products of GH action to be metabolized. Thus, genetic context can be a powerful ally in predicting the function of uncharacterized GHs.

The sulfosugar sulfoquinovose (6-deoxy-6-sulfo-glucose; SQ) is the sugar head group of the ubiquitous plant sulfolipid α-sulfoquinovosyl diacylglyceride (SQDG). SQDG is produced by almost all photosynthetic organisms and occurs in photosynthetic membranes where it supports the function of photosynthetic proteins.^3, 4^ GH family 31 (GH31) is a large glycoside hydrolase family with >20,000 members that possess diverse activities on α-glycosidic bonds and contains α-glucosidases, α-glucan lyases, α-xylanases and sulfoquinovosidasess (SQases).^5^ SQases catalyze hydrolysis of glycosides of SQ,^6, 7^ such as in SQDG and its delipidated form, sulfoquinovosyl glycerol (SQGro).^6, 7^ ^3,4^Hydrolysis releases SQ, which can be utilized as a nutrient by microorganisms through sulfoglycolysis and sulfolysis pathways.^8, 9^ Thus, SQases act as the gateway enzyme for SQ glycosides to undergo further catabolism. Family GH31 SQases are usually found within gene clusters encoding assorted sulfoglycolysis and sulfolysis pathways. Recently, a new SQ assimilation pathway, the sulfolytic sulfoquinovose monooxygenase (sulfo-SMO) pathway, was reported that involves oxidative desulfurization of SQ to release sulfite. Representative sulfo-SMO gene clusters in *Agrobacterium tumefaciens* and *Novosphingobium aromaticivorans* encode family GH31 SQases (**Fig. 1b**).^10, 11^

**Fig. 1.**
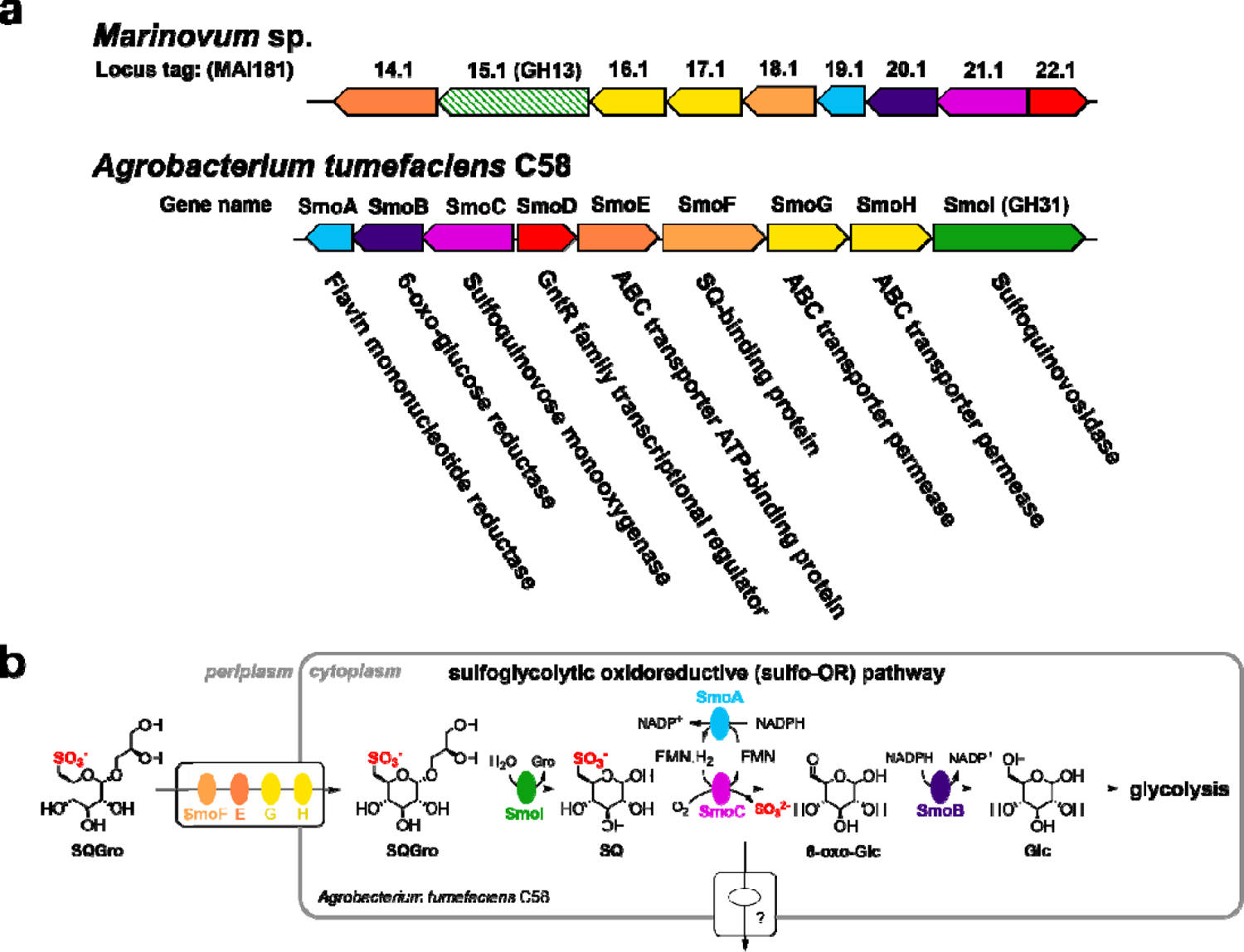
*Marinovum* sp. contains a putative sulfolytic sulfoquinovose monooxygenase (sulfo-SMO) gene cluster but lacks a candidate sulfoquinovosidase (SQase). a) Sulfolytic sulfoquinovose monooxygenase gene clusters from *Marinovum* sp. (MAI181XX.1) and *A. tumefaciens* (also known as *Agrobacterium fabrum*) strain C58. The gene coloured solid green encodes an SQase from glycoside hydrolase (GH) family 31; the gene coloured striped green encodes a member of family GH13. The *Marinovum* sp. cluster was identified using MultiGeneBLAST (see Material and Methods). **b)** The sulfo-SMO pathway of *A. tumefaciens* for desulfurization of sulfoquinovosyl glycerol (SQGro). SQ = sulfoquinovose, Glc = glucose.

By searching using the MultiGeneBLAST tool,^12^ we discovered a *Marinovum* sp. (a member of the *Roseobacter* clade bacteria)^13^ that contains a cluster of genes encoding proteins that are homologous to the key sulfo-SMO enzymes, suggesting that it encodes a sulfo-SMO pathway (**Fig. 1a**). However, this proposed gene cluster lacked a candidate gene encoding a family GH31 SQase. Instead, the gene cluster contained a gene encoding a GH13 protein (hereafter *Ms*GH13). Family GH13 is the largest of the CAZy families of glycoside hydrolases, and contains approximately 160,000 sequences but with fewer than 800 members that have been biochemically characterized to define their substrate.^14^ Family GH13 includes enzymes with activities on α-glycosidic bonds including α-glucosidases, α-amylases, α-glucosyltransferases, and sucrose α-glucosidases. Thus, families GH13 and GH31 share some overlapping activities. However, no SQases have been described in family GH13.

In this work we study the substrate specificity of *Ms*GH13. We clone and express *Ms*GH13 and screen it against a panel of activated, 4-nitrophenyl α-glycosides. We show that *Ms*GH13 lacks SQase activity, but is an α-glucosidase active on PNPGlc. Further analysis reveals it to be active on a wide range of different α-glucosidic linkages, namely maltose, sucrose, nigerose, trehalose, isomaltose and kojibiose. We use AlphaFold to present a model for the *Ms*GH13 enzyme and its active site.

## Materials and Methods

### Bioinformatic identification of *Marinovum* sp. sulfo-SMO locus

The *A. tumefaciens* SQase (AAK90105.1, locus tag Atu3285), ABC transporter membrane spanning domains (AAK90106.2 and AAK90107.2, locus tags Atu3284 and Atu3283), ABC transporter substrate binding domain (AAK90108.1, locus tag Atu3282), ABC transporter ATP binding domain (AAK90109.2, locus tag Atu3281), sulfo-SMO transcriptional regulator (AAK90110.1, locus tag Atu3280), SQ monooxygenase (AAK90111.1, locus tag Atu3279), 6-oxo-Glc reductase (AAK90112.1, locus tag Atu3278) and flavin reductase (AAK90113.1, locus tag Atu3277) were submitted separately as queries to the NCBI protein-protein BLAST^15^ (BLASTp) (https://blast.ncbi.nlm.nih.gov/). The database used was non-redundant protein sequences (nr), with *A. tumefaciens* (taxid: 358) sequences excluded. Standard algorithm parameters were used, except the maximum target sequences was set to 10,000. The raw BLASTp results were filtered by *E*-value threshold, with protein sequences retained if *E*-value ≤ 1.19e-51, the smallest of the maximum *E*-values obtained from the BLASTp searches of the three enzymes that interact with the SQ moiety (sulfoquinovosidase, SQ monooxygenase and the ABC transporter substrate binding domain). The lists of protein sequences were converted to lists of the corresponding nucleotide sequences. Protein sequences that did not return a corresponding nucleotide sequence were discarded. The nine lists were combined, with duplicates and any remaining *A. tumefaciens* sequences removed to give a list of 858,315 candidate nucleotide sequences. Python scripts were used to filter BLASTp results by *E*-value threshold, convert protein sequences to corresponding nucleotide sequences, combine the lists, remove duplicates, and remove any *A. tumefaciens* sequences. These scripts are available on GitHub, along with instructions for their use (https://github.com/jmui-unimelb/Gene-Cluster-Search-Pipeline).^16^

The list of candidate sequences was converted to a MultiGeneBLAST^17^ reference library, and searched with the *A. tumefaciens* sulfo-SMO gene cluster as a query (http://multigeneblast.sourceforge.net/). The search was run on a high-performance computing (HPC) cluster to enable parallelization (https://www.bio21.unimelb.edu.au/hpc/index.php). A modified MultiGeneBLAST script is available on GitHub that can be run in parallel on an HPC (https://github.com/jmui-unimelb/Gene-Cluster-Search-Pipeline).^16^

Data were filtered by MultiGeneBLAST score (minimum score 3.0). The MultiGeneBLAST score for a gene cluster is determined by the following equation: score = *h* + *i* × *s*. *h* is the number of query genes that had homologs (i.e., had a BLAST hit of at least a user-specified sequence coverage and % identity to the query gene; default values used in this study were sequence coverage = 25 and % identity = 30). *s* is the number of contiguous gene pairs with conserved synteny (i.e., gene order), while *i* is a factor that determines the weight of synteny in the score (default value: 0.5; default used in this study). A value of 0.5 means that the number of homologous genes is given twice the weight as conservation of gene order.^17^ Putative sulfo-SMO gene clusters identified using this workflow were screened for clusters that contained a GH13 enzyme as a candidate SQase.

### Cloning, expression and purification of *Ms*GH13

A dsDNA oligonucleotide encoding *Ms*GH13 (MAI18115.1) was synthesized (Integrated DNA Technologies) and cloned into the pET29b(+) vector at the *NdeI* and *XhoI* sites (**Fig. S1**). The resulting plasmid was sequence-verified by Sanger sequencing, then transformed into ‘T7 Express’ *E. coli* (New England BioLabs) and selected on LB-agar + 50 μg/ml kanamycin at 37 °C for 16 h. A single colony was used to inoculate 10 ml of LB media + 50 µg/ml kanamycin and the cultures were incubated at 37 °C for 16 h. These starter cultures were used to inoculate 1000 ml of S-broth (35 g tryptone, 20 g yeast extract, 5 g NaCl, pH 7.4) + 50 µg/ml kanamycin, which was incubated with shaking (250 rpm) at 37 °C until it reached an A_600_ of 0.7. The culture was cooled to room temperature, isopropyl thio-β-D-galactoside added to a final concentration of 400 µM, and incubation with shaking (200 rpm) continued at 18 °C for 19 h. Cells were harvested by centrifugation at 8,000 *g* for 20 min at 4 °C, then the pellet was resuspended in 40 ml binding buffer (50 mM NaP_i_, 300 mM NaCl, 5 mM imidazole, pH 7.5) containing protease inhibitor (Roche cOmplete EDTA-free protease inhibitor cocktail) and lysozyme (0.1 mg/ml) by nutating at 4 °C for 30 min. Benzonase (1 µl) was added to the mixture and lysis was effected by sonication [10 × (15 s on; 45 s off) at 45% amplitude]. The lysate was centrifuged at 18,000 *g* for 20 min at 4 °C and the supernatant was collected. The supernatants were filtered (0.45 µm) and loaded onto a 1 ml HisTrap IMAC column (GE). The column was washed with 10 ml of binding buffer, then the protein was eluted using elution buffer (50 mM NaP_i_, 300 mM NaCl, 400 mM imidazole, pH 7.5). Fractions containing product, as judged by SDS-PAGE, were further purified by size exclusion chromatography on a HiPrep 16/60 Sephacryl S-200 HR column (GE) using 50 mM NaP_i_, 150 mM NaCl, pH 7.5.

### Substrate screening of *Ms*GH13

Hydrolytic activity was examined towards a range of 4-nitrophenyl α-glycosides (each 4 mM) in 600 μL reaction mixtures containing 50 nM *Ms*GH13 in 50 mM citric acid/phosphate buffer, 150 mM NaCl, pH 7 at room temperature. Reactions were performed in a cuvette and monitored for release of 4-nitrophenol/4-nitrophenolate using an UV/Vis spectrophotometer at the isosbestic point (λ = 348 nm, ε = 5125 M^-1^ cm^-1^). The following sugars were examined: α-D-glucopyranoside (PNPGlc), 4-nitrophenyl α-D-sulfoquinovoside (PNPSQ), 4-nitrophenyl α-D-galactopyranoside (PNPGal), 4-nitrophenyl α-D-xylopyranoside (PNPXyl), 4-nitrophenyl α-D-mannopyranoside (PNPMan), 4-nitrophenyl *N*-acetyl-α-D-galactosaminide (PNPGalNAc), 4-nitrophenyl α-D-glucuronide (PNPGlcA) and 4-nitrophenyl α-D-glucopyranoside 6-phosphate (6-P-PNPGlc) (**Fig. S2**). PNPSQ^18^ and 6-P-PNPGlc^19^ were synthesized herein; the other PNP substrates were purchased from Sigma-Aldrich.

Hydrolytic activities toward various disaccharides were assessed in 600 μL reaction mixtures containing 73.8 nM *Ms*GH13 in 50 mM citric acid/phosphate buffer, 150 mM NaCl, pH 7 at room temperature. After 5 h, reactions were heat inactivated then applied to an aluminium backed high performance thin layer chromatography (HPTLC) sheet (Supelco) and eluted with ethyl acetate, methanol and water (7:4:2). Products were visualized by spraying with a solution of 0.2% orcinol, 10% H_2_SO_4_ and 10% water in 80% ethanol, and heating. The following sugars were assessed as substrates: trehalose [α-D-Glc*p*-(1↔1)-α-D-Glc*p*], kojibiose [α-D-Glc*p*-(1→2)-D-Glc], nigerose [α-D-Glc*p*-(1→3)-D-Glc], maltose [α-D-Glc*p*-(1→4)-D-Glc], isomaltose [α-D-Glc*p*-(1→6)-D-Glc], and sucrose [β-D-Fru*f*-(2↔1)-α-D-Glc*p*] (**Fig. S3**). A control reaction was conducted where each disaccharide was incubated under identical conditions but in the absence of enzyme. In all cases no cleavage was evident.

### pH dependence of *Ms*GH13 activity

The Michaelis parameter *k*_cat_*/K*_M_ was measured for PNPGlc hydrolysis using the substrate depletion method in 50 mM citric acid/phosphate buffer, 150 mM NaCl at a range of pH values (4.0, 4.5, 5.0, 6.0, 6.5, 7.0, 7.5, 8.0, 8.3) at room temperature. A concentration of PNPGlc of 0.1 mM was used, being <*K*_M_/10. Reactions were monitored at the isosbestic point of 4-nitrophenol (λ = 348 nm, ε = 5125 M^-1^ cm^-1^). Reactions were initiated by adding *Ms*GH13 to a final concentration of 50 nM and the rate measured continuously using a UV/visible spectrophotometer. *k*_cat_*/K*_M_ values were calculated using the equation y = (y_0_ - y_∞_)×exp(-*k*×t) + y_∞_, where *k*_cat_/*K*_M_ = *k*/[E]. p*K*_a_ values were fit to a bell shaped curve using the Prism 9.50 software package (Graphpad Scientific Software), with the equation y = m×(1/(1+[(10^-x^)/(10^-p*K*a^^1^)+(10^-p*K*a^^2^)/(10^-x^)]))+c. Below pH 4.0 and above pH 8.0 the enzyme was unstable.

### Temperature stability

Temperature stability of *Ms*GH13 was assessed by incubation of 50 nM *Ms*GH13 in the assay buffer for 3 h at different temperatures (room temperature, 30, 35, 40, 45, 50, and 55 °C). After this time, PNPGlc was added to a final concentration of 0.1 mM, and the reaction rate measured using a UV/Vis spectrometer.

### Michaelis-Menten kinetics

Michaelis-Menten kinetics were measured for *Ms*GH13 catalyzed hydrolysis of PNPGlc using a UV/visible spectrophotometer. Release of the chromogenic 4-nitrophenolate was monitored at the isosbestic point (λ = 348 nm, ε = 5125 M^-1^ cm^-1^). Reactions were conducted in 50 mM citric acid/phosphate, 150 mM NaCl (pH 6.5) at 25 °C using 50 nM *Ms*GH13 at substrate concentrations ranging from 0.1 to 2 mM.

For quantitative analysis of *Ms*GH13 hydrolysis of disaccharides, the reaction was measured by using the Colorimetric Detection Kit (Invitrogen^TM^ by Thermo Fisher Scientific). The reactions were performed in a total volume of 40 μl in an 0.5 ml Eppendorf tube by addition of 12.3 μM *Ms*GH13 to 50 mM citric acid/phosphate, 150 mM NaCl buffer (pH 6.5) with various concentrations of the disaccharide substrates. Samples at each concentration were measured after 10 and 20 min incubation to ensure a linear initial rate for the reaction. The reaction was quenched by heating samples at 80 °C for 5 minutes. The quenched samples were cooled to room temperature then glucose release was quantified according to the manufacturer’s instructions at 30 °C in a a 96-well plate using a UV/visible spectrophotometer. Hydrolysis of maltose, nigerose, trehalose, kojibiose and isomaltose each produce two equivalents of glucose, and the reaction rates were therefore halved. Control reactions were conducted under identical conditions but without addition of *Ms*GH13, and resulted in no detectable glucose release for all disaccharides. Kinetic parameters (*k*_cat_, *K*_M_, *k*_cat_*/K*_M_) were calculated using the Prism 9.50 software package (GraphPad Scientific Software) using the Michaelis-Menten equation.

### NMR spectroscopy

The stereochemistry of the initially formed product of *Ms*GH13 catalyzed hydrolysis of PNPGlc, was monitored by ^1^H NMR spectroscopy (500 MHz). *Ms*GH13 (73.8 nM) was added to a solution of PNPGlc (10 mM) in 50 mM citric acid/phosphate buffer, 150 mM NaCl at pD 7.4.

### 3D modelling

A 3D model for *Ms*GH13 was constructed using Alphafold.^20, 21^ An overlay with *Halomonas* sp. H11 glucosidase (PDB 3WY4) was generated using the *align* function within PyMOL.

## Results

MultiGeneBLAST was used to search the non-redundant protein sequence (nr) database for organisms that possess groups of genes encoding proteins of the sulfoglycolytic monooxygenase pathway: SmoA, SmoB, SmoC, SmoD and SmoI. Putative sulfo-SMO gene clusters were filtered and manually screened for the absence of a homolog of a gene encoding the family GH31 SmoG SQase, leading to the identification of *Marinovum* sp. (GenBank: NZGN01000053.1), containing a gene encoding a family GH13 protein, *Ms*GH13.

Recombinantly expressed *Ms*GH13 (MAI18115.1) was screened against various PNP glycosides: PNPGlc was a good substrate, but it was inactive against PNPSQ, PNPGal, PNPXyl, PNPMan, PNPGalNAc, PNPGlcA and 6-P-PNPGlc (for structures see **Fig. S2**).

The stereochemical outcome of *Ms*GH31 catalyzed hydrolysis of PNPGlc was assessed by ^1^H NMR spectroscopy. This revealed the initial formation of α-glucose (δ 5.15 ppm, d, *J*_1,2_ 3.8 Hz), which slowly underwent mutarotation to give a 1:1.42 ratio of α-glucose and β-glucose (δ 4.56 ppm, d, *J*_1,2_ 8.0 Hz). This demonstrated that the enzyme is a retaining a α-glucoside hydrolase (**Fig. 2**).

**Fig. 2.**
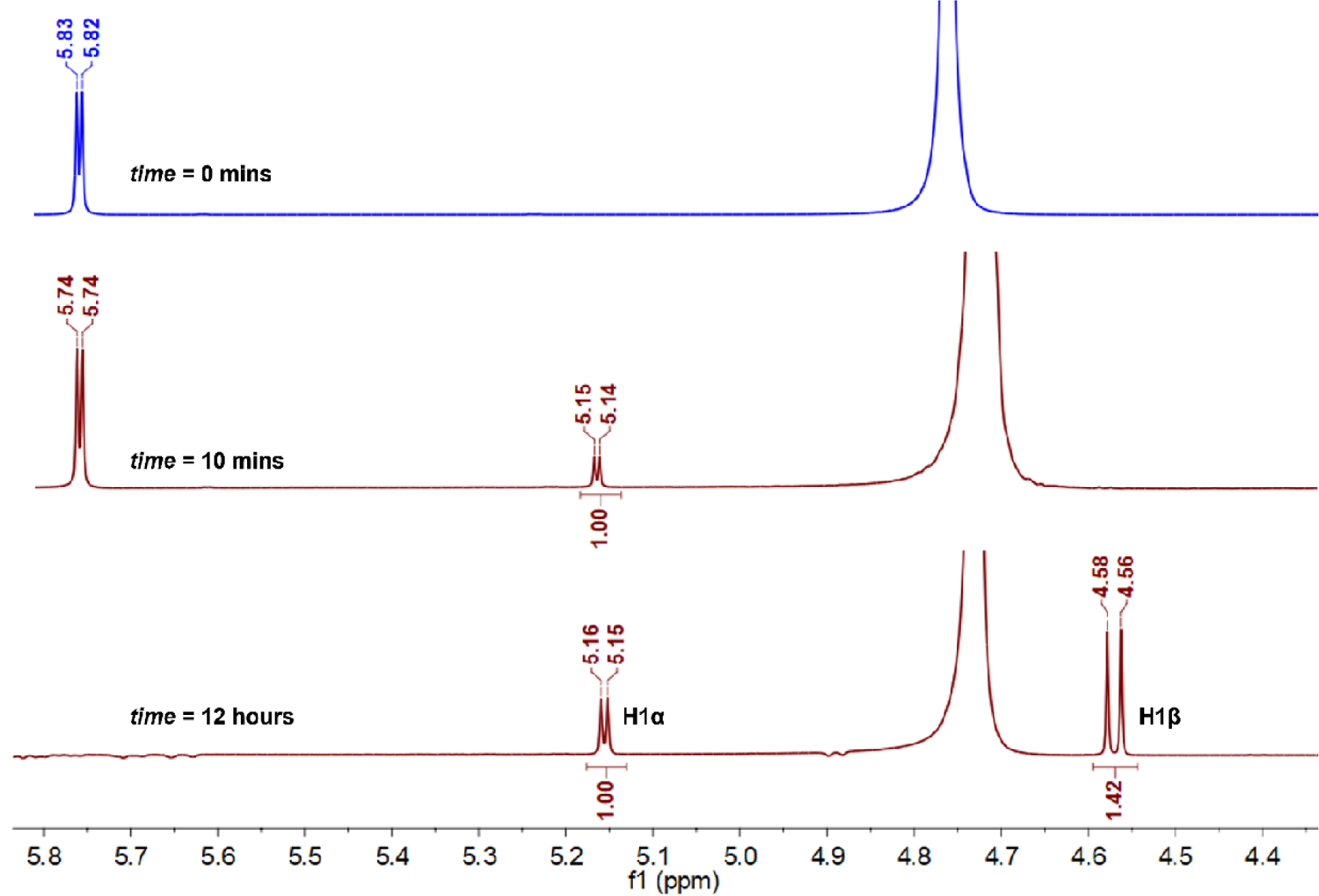
*Ms*GH13 is a stereochemically retaining α-glucosidase. Excerpt of ^1^H-NMR spectra (500 MHz, D_2_O) of 4-nitrophenyl α-glucopyranoside (PNPGlc) (top) and the reaction course of *Ms*GH13 catalyzed hydrolysis of PNPGlc at 10 min and 12 h. H1 of PNPGlc is present at δ 5.82 ppm. At *t* = 10 min, the initially formed signal at δ 5.15 ppm, *J*_1,2_ 3.8 Hz, is assigned as H1 of α-glucose (middle). At *t* = 12 hours the α-glucose underwent mutarotation and formed a signal at δ 4.6 ppm, *J*_1,2_ 8.0 Hz, assigned H1 of β-glucose (bottom).

The effects of pH on the hydrolytic activity and the temperature stability of *Ms*GH13 were evaluated using PNPGlc as a substrate. The pH dependence of *k*_cat_/*K*_M_ gave a bell-shaped curve, with an optimum at pH 6.5, and p*K*_a1_ and p*K*_a2_ values of 6.2 ± 0.2 and 7.1 ± 0.2, respectively (**Fig. 3a**). The effect of temperature on *Ms*GH13 activity was assessed at pH 6.5. *Ms*GH13 incubated at 25 °C retained activity, enzyme incubated at 30 °C retained only 16.7% activity, and enzyme incubated at 35 °C and above lost all activity. Kinetic parameters for PNPGlc measured at pH 6.5 and 25 °C were *k*_cat_ = 12.4 s^-1^, *K*_M_ = 0.95 mM and *k*_cat_*/K*_M_ = 13,000 M^-1^ s^-1^ (**Fig. 3b**).

**Fig. 3.**
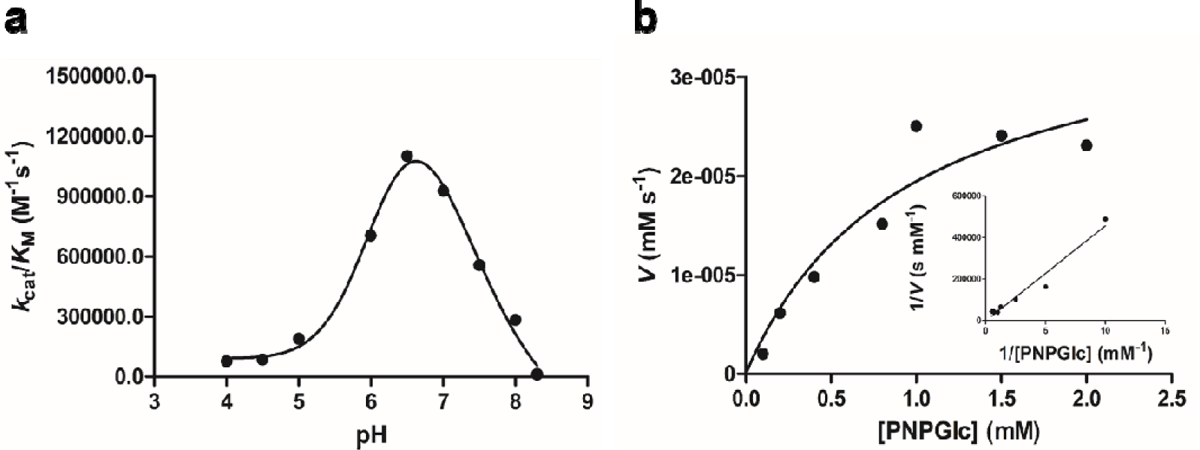
*Ms*GH13 from *Marinovum* sp. is an α-glucosidase active on 4-nitrophenyl α-glucopyranoside (PNPGlc). **a)** pH dependence of *k*_cat_/*K*_M_ using PNPGlc as substrate. **b)** Michaelis-Menten plot for PNPGlc as substrate. (inset) Lineweaver-Burk plot.

Hydrolytic activity on trehalose, kojibiose, nigerose, maltose, isomaltose, and sucrose was qualitatively assessed by HPTLC. Nigerose, trehalose, sucrose and maltose were completely hydrolyzed, while kojibiose and isomaltose were partially hydrolyzed (**Fig. 4a**). Michaleis-Menten kinetic parameters reveal the greatest activity for maltose (*k*_cat_ = 135 s^-1^, *K*_M_ = 3.22 mM and *k*_cat_*/K*_M_ = 42,000 M^-1^ s^-1^) (**Table 1**). The relative percentage of *k*_cat_/*K*_M_ values for the other disaccharides versus maltose were: nigerose (52%), trehalose (20%), sucrose (7.9%), isomaltose (6.4%) and kojibiose (3.8%).

**Fig. 4.**
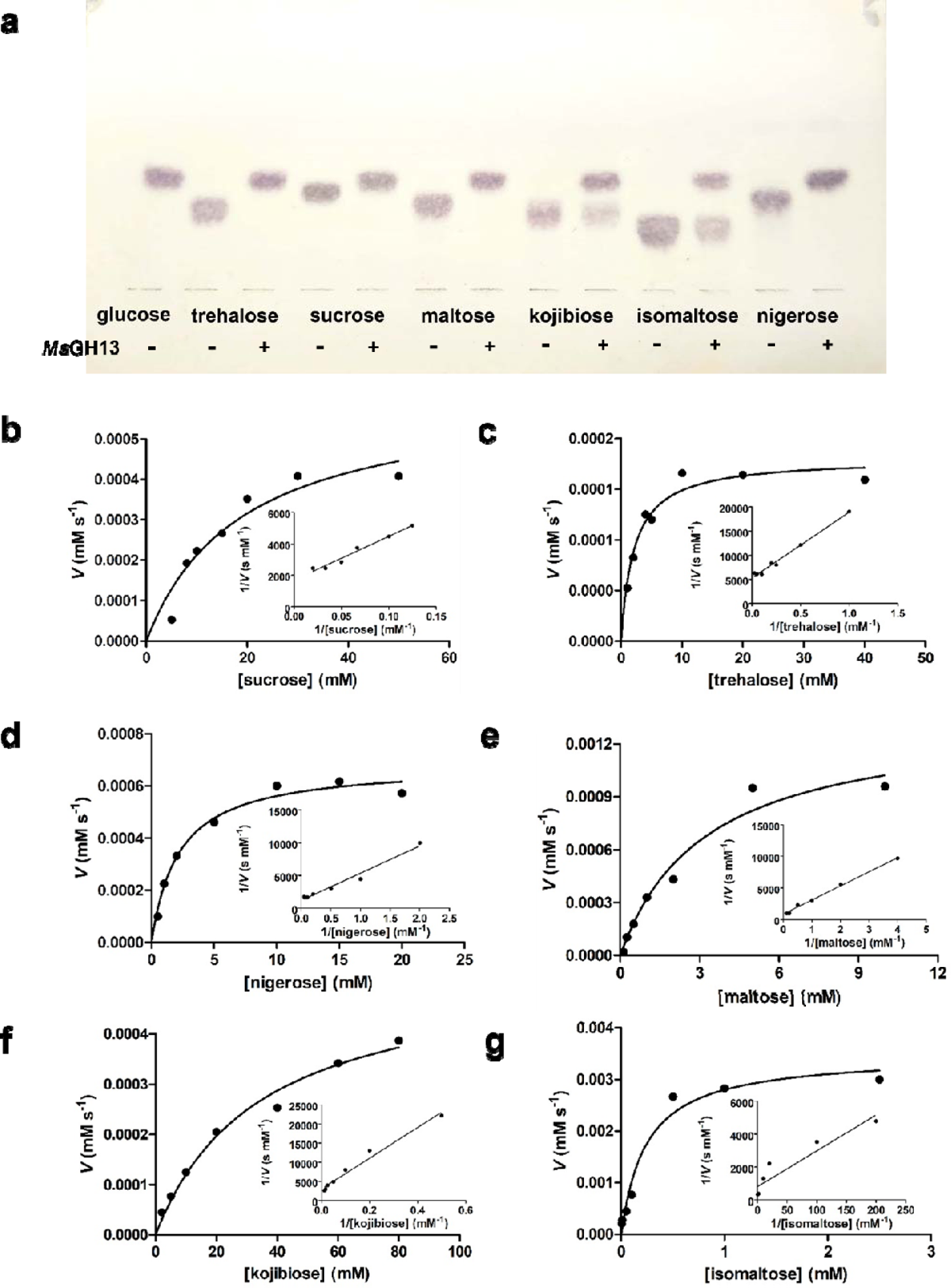
*Ms*GH13 from *Marinovum* sp. is an α-glucosidase with activity on sucrose, trehalose, nigerose, maltose, kojibiose and isomaltose. **a)** High performance thin layer chromatography analysis of *Ms*GH13 digests of various disaccharides. Products were visualized with 0.2% orcinol in 10% H_2_SO_4_, 10% H_2_O in ethanol. Michaelis-Menten and (inset) Lineweaver-Burk plots for **b)** sucrose, **c)** trehalose, **d)** nigerose, **e)** maltose, **f)** kojibiose and **g)** isomaltose.

**Table 1:** Michaelis-Menten kinetic parameters for *Ms*GH13 from *Marinovum* sp.

| Substrate | $k_{\text{cat}}$ ($\text{s}^{-1}$) | $K_{\text{M}}$ (mM) | $k_{\text{cat}}/K_{\text{M}}$<br>($\text{M}^{-1} \text{s}^{-1}$) $\times 10^3$ | Relative<br>activity,<br>($k_{\text{cat}}/K_{\text{M}}$) % |
| --- | --- | --- | --- | --- |
| PNPSQ | nd | nd | nd | 0 |
| PNPGlc | 12.4±3.3 | 0.95±0.47 | 13±7.3 | 31 |
| trehalose | 18.0±0.9 | 2.13±0.43 | 8.5±1.8 | 20 |
| kojibiose | 53.2±5.1 | 34.2±7.5 | 1.6±0.4 | 3.8 |
| nigerose | 68.3±3 | 3.1±0.36 | 22±2.7 | 52 |
| maltose | 135±17 | 3.22±0.99 | 42±14 | 100 |
| isomaltose | 0.66±0.03 | 0.24±0.07 | 2.7±0.8 | 6.4 |
| sucrose | 61.2±10.6 | 18.8±7.1 | 3.3±1.4 | 7.9 |

nd = no activity was detected

## Discussion

We used MultiGeneBLAST to identify *Marinovum* sp., which possessed genes encoding homologues of the *Agrobacterium tumefaciens* sulfoquinovose monooxygenase SmoC (75% identity), flavin reductase SmoA (47% identity), 6-oxo-glucose dehydrogenase SmoB (61% identity), as well as an ABC transporter cassette and an associated substrate binding protein. The *Marinovum* sp. gene cluster contained a gene encoding a member of family GH13, and which was computer annotated as an α-glucosidase. Family GH13 has been divided into 35 subfamilies based on sequence similarity, structure and function.^14^ The protein from *Marinovum* sp. belongs to the GH13_23 subfamily, which contains >3200 members, but just 9 members of this subfamily have been characterized. Within subfamily GH13_23, these activities include α-glucosyltransferase (EC 2.4.1.-), oligo-α-1,6-glucosidase (EC 3.2.1.10, including isomaltose), exo-1,3-α-glucanase (3.2.1.84, including nigerose), α-glucosidase (EC 3.2.1.20, including maltose and sucrose), kojibiose hydrolase (EC 3.2.1.216, including kojibiose), and α,α-trehalase (EC 3.2.1.28) activities.^22–25^

Based on the genomic context of the annotated GH13 gene in *Marinovum* sp., we set out to explore whether this might encode an SQase or another enzyme activity. *Ms*GH13 was active upon PNPGlc but was inactive against other α-glycosides including PNPSQ, PNPGal, PNPXyl, PNPMan, PNPGalNAc, PNPGlcA and 6-P-PNPGlc. Thus, *Ms*GH13 is an α-glucosidase, and is not an SQase. ^1^H NMR analysis of the products from *Ms*GH13 cleavage of PNPGlc revealed initial formation of α-Glc, which mutarotated to give an equilibrium mixture of α- and β-glucose; thus, *Ms*GH13 is a stereochemically retaining α-glucosidase.

*Ms*GH13 catalyzed the hydrolysis of the following disaccharides: trehalose, kojibiose, nigerose, maltose, isomaltose, and sucrose. The second order rate constants *k*_cat_/*K*_M_ revealed a 25-fold range of activity, with a 2-fold preference for maltose over nigerose, and greater ratios for the remaining disaccharide substrates (**Fig. 4b-4g, Table 1**). Thus, *Ms*GH13 is an unusually broad-spectrum α-glucosidase with the ability to cleave α-1,1-, 1,2-, 1,3-, 1,4- and 1,6-glucosidic linkages. This ability to cleave diverse linkages is shared by *Thermus thermophilus* HB27 α,α-trehalase, another member of subfamily GH13_23. While HB27 prefers trehalose, it displays good relative activity on isomaltose (63%) and kojibiose (20%), and modest activities on nigerose (16%), maltose (10%) and sucrose (8%).^25^

Comparison of the AlphaFold 3D structure prediction of *Ms*GH13 to the experimentally determined 3D X-ray structure of *Halomonas* sp. H11 glucosidase (HaG) mutant with maltose^26^ (PDB: 3WY4, green), which is also a member of subfamily GH13_23, shows excellent structural similarity with RMSD = 2.24 Å (**Fig. 5**). This overlay reveals conservation of the key catalytic machinery, allowing assignment of catalytic nucleophile (*Ms*GH13/HaG: D201/D202), acid/base (E272/E271), and substrate-binding residues (D61/D62, H104/H105, R199/R200, H336/H332, D337/D333 and R410/R400). Like *Ms*GH13, HaG prefers maltose (100%).^27^ However, HaG is much more specific, with good relative activity on sucrose (33%), very low relative hydrolysis activity against isomaltose (1.1%), trehalose (2.6%), nigerose (4.2%), and no detectable activity on kojibiose.^27^ These active site residues are also conserved within the broad-spectrum HB27 α,α-trehalase.^25^

**Fig. 5.**
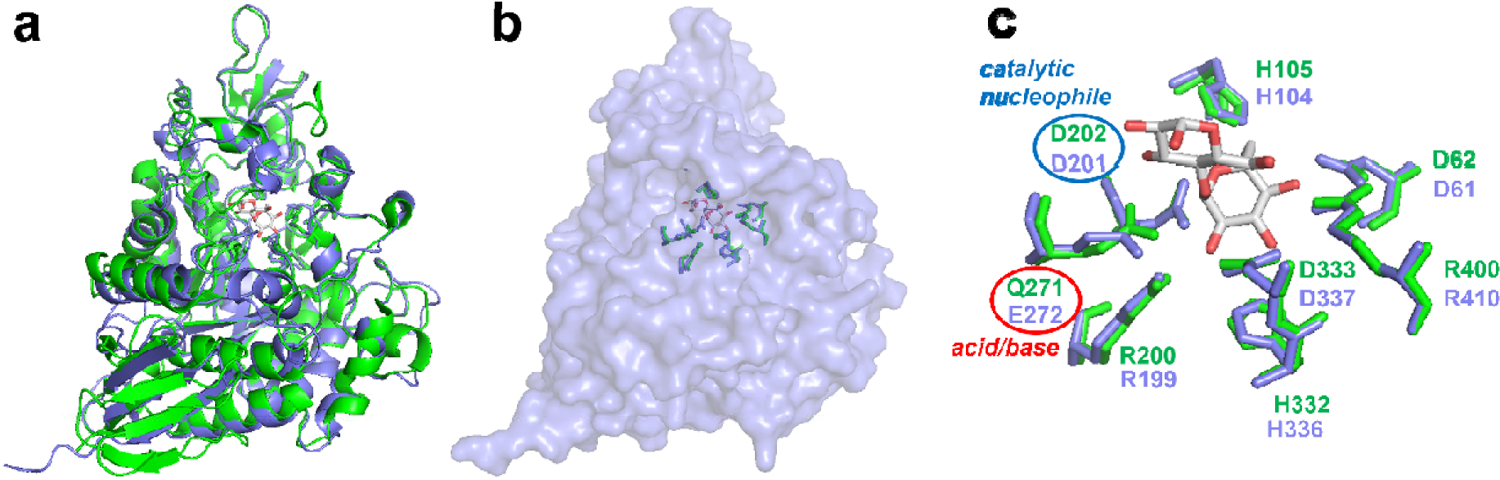
*Ms*GH13 is predicted to recognize substrate in a similar way to the more substrate specific GH13 member *Halomonas* sp. H11 glucosidase (HaG). **a)** Overlay of *Ms*GH13 (AlphaFold predicted 3D structure, blue) and HaG E271Q mutant with maltose (PDB: 3WY4, green).^26^ **b)** Surface presentation of *Ms*GH13 predicted structure with active site residues and overlay of maltose from HaG complex. **c)** Comparison of overlay of Alphafold prediction of *Ms*GH13 with Michaelis complex of HaG with maltose. The overlay reveals conservation of the key catalytic machinery, allowing assignment of catalytic nucleophile (*Ms*GH13/HaG: D201/D202), acid/base (E272/Q271 (corresponding to E271), respectively), and substrate-binding residues (D61/D62, H104/H105, R199/R200, H336/H332, D337/D333 and R410/R400, respectively).

This study was initiated based on the genomic context of *Ms*GH13 within a putative sulfo-SMO gene cluster, and the observation that sulfoglycolytic and sulfolytic gene clusters typically contain SQase genes of GH family 31. *Marinovum* sp. is a member of the *Roseobacter* clade bacteria and a recent study by Tang *et al.* demonstrated that other Roseobacter clade bacteria containing a sulfo-SMO gene cluster, *Dinoroseobacter shibae* DFL 12 and *Roseobacter denitrificans* OCh 114, can grow on SQ.^28^ These two bacteria and many more *Roseobacter* clade bacterial genomes that contained putative sulfo-SMO metabolic genes (including *Marinovum algicola* DG 898) also lack a family GH31 SQase candidate.^28^ Tang *et al.* reported that marine phytoplankton and cyanobacteria contain intracellular SQ at micro to millimolar levels, and that SQ can be detected within particulate organic matter sampled from the Bohai Sea, suggesting that *Roseobacter* clade bacteria may encounter sufficient environmental SQ to support their growth without a requirement for an SQase.^28^

In conclusion, *Ms*GH13, while encoded within a genomic context that is suggestive of possible SQase activity, is really a broad spectrum α-glucosidase with activity on a wide range of α-linked disaccharides. The high similarity of the active sites of the AlphaFold predicted *Ms*GH13 structure with that of the maltose/sucrose specific HaG α-glucosidase provides little insight into how the active site of the former can accommodate such a broad range of disaccharides. Possibly, this reflects limitations in the use of AlphaFold to predict structures of complexes, and provides an impetus for structural studies to define in greater detail how disaccharides with other linkages can be accommodated within the active site.

## Declarations

**Ethical Approval** Not applicable

**Consent to participate** Not applicable

**Consent to publish** Not applicable

## Author contributions

SJW conceived the study with input from EGB and DA. JL, JM, BMdS, DEVP and DBA performed bioinformatics. NMS and EGB conducted molecular biology. JL conducted enzymology. JL and SJW wrote the paper with input and approval of the final draft from the authors.

## Funding

S.J.W is supported by the Australian Research Council (DP210100233, DP210100235). D.B.A. is supported by an Investigator Grant from the National Health and Medical Research Council (NHMRC) of Australia (GNT1174405). E.D.G-B. acknowledges support from The Walter and Eliza Hall Institute of Medical Research, National Health and Medical Research Council of Australia (NHMRC) Ideas grant GNT2000517, the Australian Cancer Research Fund; and the Brian M. Davis Charitable Foundation Centenary Fellowship. JL thanks the China Scholarship Council for a PhD scholarship. BMdS is supported by Melbourne Research Scholarship. Phillip van der Peet is thanked for providing 6-P-PNPGlc.

## Competing interests

The authors declare that they have no conflict of interest.

## Availability of data and materials

All data generated or analysed during this study are included in this published article and its supplementary information files.

## Notes

### Competing Interest Statement

The authors have declared no competing interest.

### Summary of Updates

Title changed. A range of changes to text in response to referee comments upon review.

