## Supplementary material file for "A broad-spectrum α-glucosidase of glycoside hydrolase family 13 from *Marinovum* sp., a member of the *Roseobacter* clade"

**Synthetic dsDNA:**

GCCAGTCATATGTCCAGCCTGAAATGGTGGGAGCATGCTGTCATTTACCAGATTTATCCTCGTTCCTTTCAGGACAGCAACTCTGACGGCATCGGTGATCTGAACGGCATTACCTCCCGTCTGGAATACATCGCTAACCTGGGTGTAGACGCGATCTGGATTTCCCCGTTCTTCATGAGCCCGCAACACGACTTTGGCTACGATGTTTCCAACTACTGCGAAGTTAATCCGGAGTACGGCAGCCTGTCTGATTTCGATCAGCTGATCGACAAGGTGCACGGTCTGGGTCTGCGCCTGATGATTGACATCGTCCCGGCACATTGTTCTTACCTGCACCCGTGGTTCGAAGAAAGCCGTAAAAGCAAAGACAACCCAAAATCCGATTGGTTCCACTGGGTTGATCCTAAACCGGACGGTTGTCCGCCGAACAACTGGCTGTCCTTCTTCGGTGGTCCTGCATGGAGCTGGGAACCGCGTCGTCAACAGTACTACCTGCACAACTTCCTGCCGGAACAGCCTAACCTGAACCACGCAAACCAGGAAGTTCAGGAAGAACTGAACAAGGTAGCGCGCTTTTGGTTCGATCGTGGCGTTGATGGCTTTCGTCTGGACGCAGTACACACCGTTAATGGCGACTGCGAACCGTATAAGGACAACCTGGCCGACCCGAACTTTACTCTGGGTCCGCTGCCTCAGGATAAACAGCCGTTCTTCCGTCAGCTGCACGACGTTGGCCAGCTGAACCAGCCGGTTATTCAGAAATTTTCCGAAGCGTATCGTAAGATCGCAGATGGTTATGACGGCGACCGCTTCCTGATGGGTGAAGTTGACGGCGATGATGGTAACGCTATGGCGGTTAGCAAAACCTTTTCTGAACCGGGTCGTCTGCATAGCACTTATAACTTCGATCTGCTGGAATGGGGCGGCCTGAACGTCAGCGAACTGAAACAGGCTATCGAAAACGCAAAAGAAGTATTCAACGGCAGCGGCCGTCTGTGCTTCGCATTCAGCAACCATGATGTGCCGCGTTCTGCATCTCGTCAGCTGGACCCGCTGGGCATTACTTCTGATAAACAGGACGATCTGCAGCTGCTGCTGCTGCAGCTGGAAACCTCTCTGATCGGCTCTTCCTGTATTTATCAGGGCGAAGAACTGGGCCTGTCTGATGTCACCGATATCGAATTCGATAAGATGAAAGATCCGTGGGGTATCAACTTCTACCCGGAATTTCTGGGTCGTGACACCTGCCGTACTCCGATGGTTTGGGAAAAGAGCAAGCCGATGGGTGGTTTTACTAGCGCGAACGAATCTTGGCTGCCGATCTCCAAGTCTCACCTGGAGAAGGCGGGTCTGGACATGGCAAAATCTGAAGGTTCCATCTACAACAAATTTTCTTCTTTTCTGAAGTGGCGTAAACAGCAGCCTGCGCTGATGACTGCTAATAACATGAGCTCTATCACTGGTGGCCCGCGTGAAATCATCTTCGACCGCATCTCCAAGACTCAGACTCTGCGTTGCAAGTTCGACTTCGAACTGGTTAAAGCGACTTTTGAAGAAGTAACCCACGGTACCGGCCTCGAGCCGCTG

**CDS:**

ATGTCCAGCCTGAAATGGTGGGAGCATGCTGTCATTTACCAGATTTATCCTCGTTCCTTTCAGGACAGCAACTCTGACGGCATCGGTGATCTGAACGGCATTACCTCCCGTCTGGAATACATCGCTAACCTGGGTGTAGACGCGATCTGGATTTCCCCGTTCTTCATGAGCCCGCAACACGACTTTGGCTACGATGTTTCCAACTACTGCGAAGTTAATCCGGAGTACGGCAGCCTGTCTGATTTCGATCAGCTGATCGACAAGGTGCACGGTCTGGGTCTGCGCCTGATGATTGACATCGTCCCGGCACATTGTTCTTACCTGCACCCGTGGTTCGAAGAAAGCCGTAAAAGCAAAGACAACCCAAAATCCGATTGGTTCCACTGGGTTGATCCTAAACCGGACGGTTGTCCGCCGAACAACTGGCTGTCCTTCTTCGGTGGTCCTGCATGGAGCTGGGAACCGCGTCGTCAACAGTACTACCTGCACAACTTCCTGCCGGAACAGCCTAACCTGAACCACGCAAACCAGGAAGTTCAGGAAGAACTGAACAAGGTAGCGCGCTTTTGGTTCGATCGTGGCGTTGATGGCTTTCGTCTGGACGCAGTACACACCGTTAATGGCGACTGCGAACCGTATAAGGACAACCTGGCCGACCCGAACTTTACTCTGGGTCCGCTGCCTCAGGATAAACAGCCGTTCTTCCGTCAGCTGCACGACGTTGGCCAGCTGAACCAGCCGGTTATTCAGAAATTTTCCGAAGCGTATCGTAAGATCGCAGATGGTTATGACGGCGACCGCTTCCTGATGGGTGAAGTTGACGGCGATGATGGTAACGCTATGGCGGTTAGCAAAACCTTTTCTGAACCGGGTCGTCTGCATAGCACTTATAACTTCGATCTGCTGGAATGGGGCGGCCTGAACGTCAGCGAACTGAAACAGGCTATCGAAAACGCAAAAGAAGTATTCAACGGCAGCGGCCGTCTGTGCTTCGCATTCAGCAACCATGATGTGCCGCGTTCTGCATCTCGTCAGCTGGACCCGCTGGGCATTACTTCTGATAAACAGGACGATCTGCAGCTGCTGCTGCTGCAGCTGGAAACCTCTCTGATCGGCTCTTCCTGTATTTATCAGGGCGAAGAACTGGGCCTGTCTGATGTCACCGATATCGAATTCGATAAGATGAAAGATCCGTGGGGTATCAACTTCTACCCGGAATTTCTGGGTCGTGACACCTGCCGTACTCCGATGGTTTGGGAAAAGAGCAAGCCGATGGGTGGTTTTACTAGCGCGAACGAATCTTGGCTGCCGATCTCCAAGTCTCACCTGGAGAAGGCGGGTCTGGACATGGCAAAATCTGAAGGTTCCATCTACAACAAATTTTCTTCTTTTCTGAAGTGGCGTAAACAGCAGCCTGCGCTGATGACTGCTAATAACATGAGCTCTATCACTGGTGGCCCGCGTGAAATCATCTTCGACCGCATCTCCAAGACTCAGACTCTGCGTTGCAAGTTCGACTTCGAACTGGTTAAAGCGACTTTTGAAGAAGTAACCCACGGTACCGGCCTCGAGCACCACCACCACCACCACTGA

**>MsGH13**

MSSLKWWEHAVIYQIYPRSFQDSNSDGIGDLNGITSRLEYIANLGVDAIWISPFFMSPQHDFGYDVSNYCEVNPEYGSLSDFDQLIDKVHGLGLRLMIDIVPAHCSYLHPWFEESRKSKDNPKSDWFHWVDPKPDGCPPNNWLSFFGGPAWSWEPRRQQYYLHNFLPEQPNLNHANQEVQEELNKVARFWFDRGVDGFRLDAVHTVNGDCEPYKDNLADPNFTLGPLPQDKQPFFRQLHDVGQLNQPVIQKFSEAYRKIADGYDGDRFLMGEVDGDDGNAMAVSKTFSEPGRLHSTYNFDLLEWGGLNVSELKQAIENAKEVFNGSGRLCFAFSNHDVPRSASRQLDPLGITSDKQDDLQLLLLQLETSLIGSSCIYQGEELGLSDVTDIEFDKMKDPWGINFYPEFLGRDTCRTPMVWEKSKPMGGFTSANESWLPISKSHLEKAGLDMAKSEGSIYNKFSSFLKWRKQQPALMTANNMSSITGGPREIIFDRISKTQTLRCKFDFELVKATFEEVTHGTGLEHHHHHH*

**Figure S1.** Nucleotide and protein sequences for *Ms*GH13.

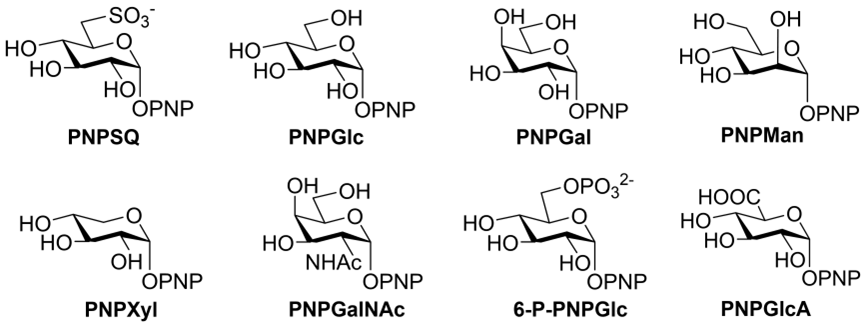

**Figure S2.** Structures of 4-nitrophenyl α-glycosides.

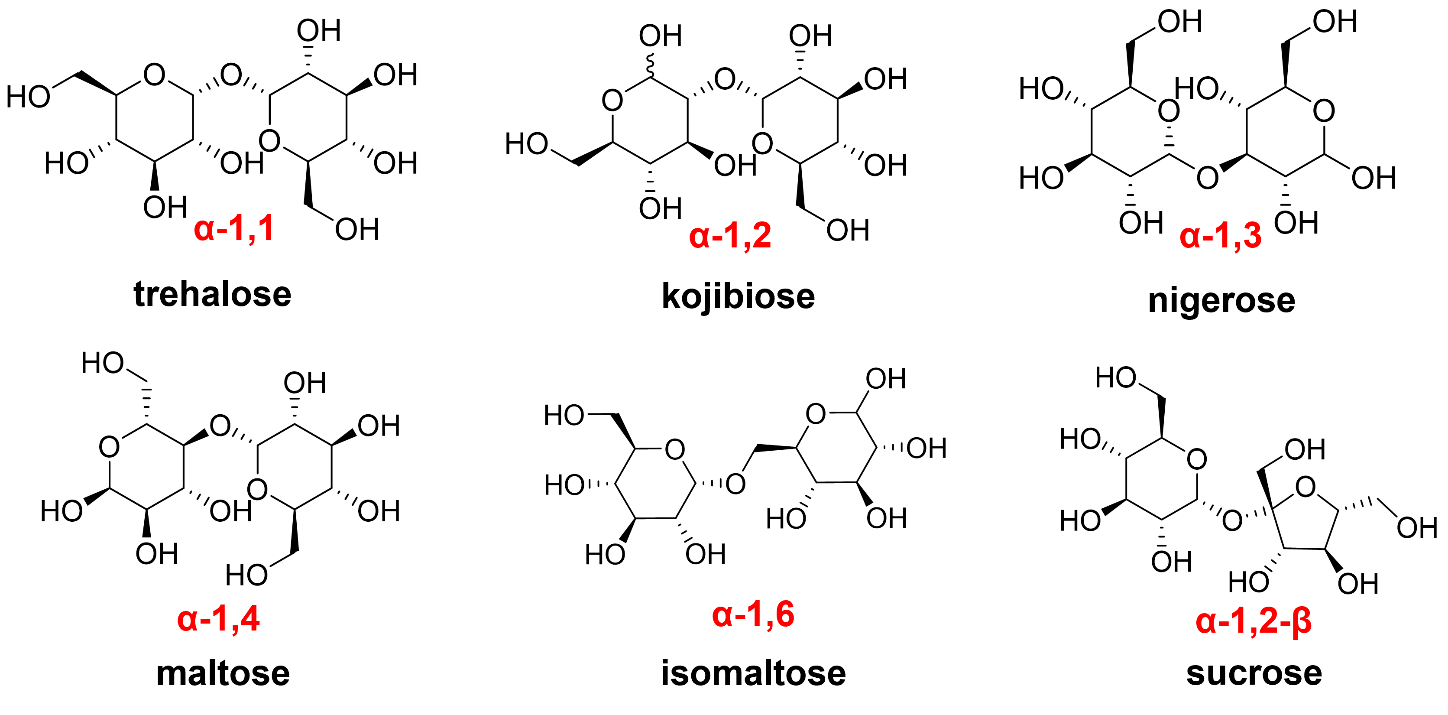

**Figure S3.** Structures of disaccharides examined as substrates.
